# Integrated valorization of glycerol and PET into lipids and PHAs using an engineered *Yarrowia lipolytica* strain and a *Pseudomonas-Comamonas* consortium

**DOI:** 10.64898/2026.06.02.729029

**Authors:** Francisco J. Molpeceres-García, Alejandro García-Miró, Alicia Prieto, David Sanz-Mata, Jorge Barriuso

**Author notes:** **Corresponding author:** Jorge Barriuso. Centro de Investigaciones Biológicas Margarita Salas (CSIC), Department of Biotechnology, Ramiro de Maeztu 9, E-28040 Madrid, Spain.

## Abstract

The accumulation of plastic waste necessitates innovative strategies that convert polymer carbon into value-added products. Here, we present a sequential yeast-bacterial workflow for the integrated valorisation of glycerol and amorphous polyethylene terephthalate (amPET) into triacyl glyceride (TAGs) and polyhydroxyalkanoates (PHAs). First, an engineered obese strain of *Yarrowia lipolytica* was cultivated on glycerol, for the simultaneous production of intracellular lipids, up to 42.9% of its cell dry weight, and secretion of a PET-depolymerizing enzymatic cocktail, composed of the cutinase from *Mycothermus thermophilus* (HiC) and the lipase B from *Moesziomyces antarcticus* (CALB). The resulting enzymatic crude hydrolysed amPET, and the released degradation products—terephthalic acid (TPA) and ethylene glycol (EG)— served as feedstocks for a bacterial consortium composed of *Comamonas testosteroni* RW31 and *Pseudomonas putida* JM37, which naturally assimilate TPA and EG, respectively. This consortium successfully upcycled the released monomers into intracellular polyhydroxybutyrate (PHB) and medium-chain-length PHAs. Furthermore, fluorescent strains of both bacteria enabled the development of a semi-quantitative method for monitoring the consortium population dynamics. Overall, this study provides a robust proof-of-concept for a circular bioeconomy approach, successfully coupling glycerol-based enzyme and lipid production with the downstream biological conversion of PET-derived monomers into valuable bioplastics.

**Graphical abstract:** 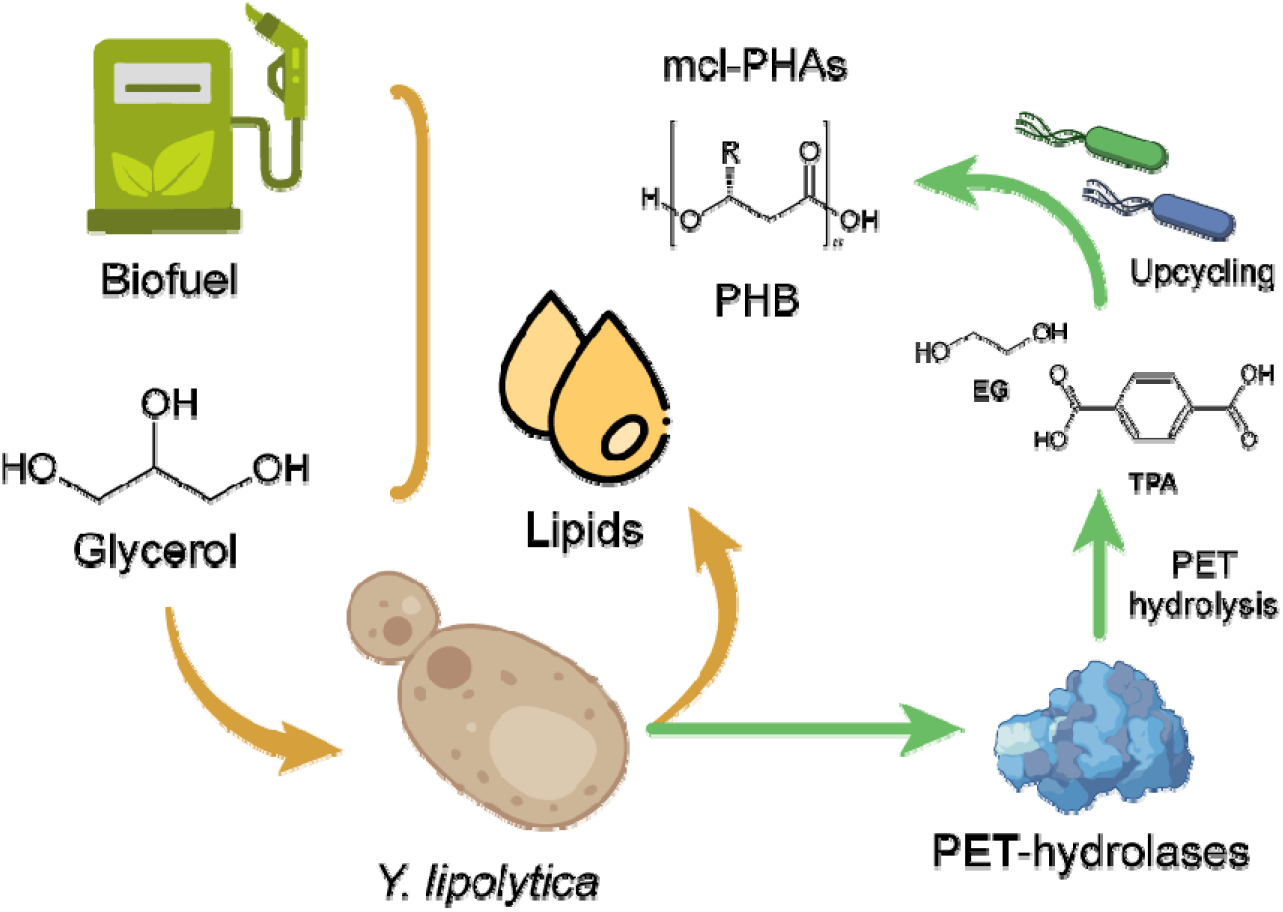

## 1. Introduction

Plastic pollution and the accumulation of post-consumer polymers have intensified the need for recycling strategies that go beyond mechanical reprocessing and enable the recovery of carbon into new value-added products. Polyethylene terephthalate (PET) is particularly relevant in this context because its ester bonds can be hydrolysed enzymatically, yielding terephthalic acid (TPA) and ethylene glycol (EG), two monomers that can subsequently be used as microbial feedstocks for the production of chemicals or materials (Diao et al. 2025; Tiso et al. 2021). This has positioned PET as a model substrate for biological recycling and upcycling.

To date, the The Plastics-Active Enzymes Database (PAZy) (Buchholz et al. 2022) contains 312 bacterial and fungal enzymes active on PET. Some of them, like the left branch compost cutinase variant LCC^ICCG^ (Arnal et al. 2023), are currently being tested at industrial scale. Despite this rapid progress in PET-active enzymes, efficient enzymatic PET hydrolysis remains limited. Crystallinity is the main bottleneck in PET hydrolysis, as efficient hydrolysis usually occurs in amorphous PET regions (Di Pede-Mattatelli et al. 2026). Hence, pre-treatments are normally required before hydrolysis. Furthermore, PET degradation generates bis(2-hydroxyethyl) terephthalate (BHET) and mono(2-hydroxyethyl) terephthalate (MHET) as intermediates, requiring complementary enzymatic activity for the subsequent conversion of MHET to TPA and EG. For this reason, combinations of polyester hydrolases and MHET/BHET-converting enzymes have been explored to increase monomer release. Among them, *Mycothermus thermophilus* (formerly *Humicola insolens*) cutinase (HiC), with activity on PET and BHET, and *Moesziomyces antarcticus* (formerly *Candida antarctica*) lipase B (CALB), with activity on MHET, have been reported as a useful enzymatic combination for PET depolymerisation, improving monomer release compared with single-enzyme reactions (Carniel et al. 2017).

The biological upcycling of PET-derived monomers requires microbial platforms able to assimilate TPA and EG and redirect their carbon flux towards products with higher value (Carniel et al. 2024). A consortium-based strategy can provide an additional advantage when different members of the community specialize in different PET-derived substrates, reducing direct competition and enabling division of labour (Bao et al. 2023). In this sense, the bacteria *Comamonas testosteroni* RW31 and *Pseudomonas putida* JM37 can naturally assimilate TPA and EG, respectively, and convert these monomers into valuable compounds such as the bioplastics polyhydroxyalkanoates (PHAs) or the pigment violacein (Molpeceres-García et al. 2025; Molpeceres□García et al. 2026). Here, developing techniques for an easy monitorization of the population dynamics would be of interest.

In parallel, the development of PET upcycling schemes should also consider how the enzymes required for depolymerisation are produced. The oleaginous yeast *Yarrowia lipolytica* is an attractive host for this purpose because it combines the assimilation of a broad range of substrates, robust growth, the availability of tools for genetic manipulation, secretion capacity, and a strong ability to accumulate intracellular lipids under nitrogen-limited conditions (Park & Ledesma-Amaro 2023). In this sense, *Y. lipolytica* can be used as a chassis to produce extracellular enzymes, including PET-hydrolases (Kosiorowska et al. 2022; Theron et al. 2020), and it can use glycerol, including glycerol derived from biodiesel production, as a low-cost carbon source for biomass and lipid production (Juszczyk et al. 2013). Therefore, a glycerol-fed *Y. lipolytica* module could potentially serve two functions within an upcycling workflow: production of extracellular PET-hydrolytic enzymes and simultaneous generation of microbial lipids as a coproduct.

Here, we investigated a sequential yeast–bacterial fermentation strategy for the coupled valorisation of glycerol and amorphous PET. An engineered obese strain of *Y. lipolytica* was used to convert glycerol into intracellular lipids while secreting an enzymatic cocktail composed of HiC and CALB. The resulting crude enzyme preparation was then applied for amorphous PET hydrolysis, and the released PET-derived monomers were used as feedstock for a *C. testosteroni* RW31–*P. putida* JM37 consortium able to accumulate PHAs. In addition, fluorescently labelled derivatives of both bacterial strains were constructed to facilitate monitoring of consortium dynamics. Overall, this work provides a proof of concept for connecting glycerol-based enzyme/lipid production with downstream biological conversion of PET-derived monomers into PHAs.

## 2. Materials and Methods

### 2.1 Design and construction of recombinant strains

The obese strain DGA of *Y. lipolytica* was previously constructed (De La Torre et al. 2026) using *Y. lipolytica* Po1d as background strain (Niehus et al. 2018). In brief, codon-optimized genes *dga1* from *Rhodotorula toruloides* (formerly *Rhodosporidium toruloides*) and *dga2* from *Claviceps purpurea* were cloned under a constitutive pTEF promoter into the *mfe1* locus using the Addgene Golden Gate kit for *Y. lipolytica* (ref. 1000000167) (Larroude et al. 2019), in order to promote triacyl glyceride (TAGs) production and limit fatty-acid β-oxidation (Friedlander et al. 2016; Yan et al. 2020). To express and secrete active PET-hydrolases in *Y. lipolytica* DGA, codon-optimized *hic* (*Mycothermus thermophilus* cutinase) and *calb* (*Moesziomyces antarcticus* Lipase B) genes, each fused with a SP3 signal peptide (Celińska et al. 2020), were cloned into the JMP62Leu plasmid (Nicaud et al. 2002) and transformed into *Y. lipolytica* DGA, obtaining the strain *Y. lipolytica* HC (García-Miró et al. 2026).

In the case of the bacteria, *C. testosteroni* RW31 and *P. putida* JM37, there were transformed with fluorescent proteins to easily track their population dynamics in co-culture. A constitutive *Ptrc* promoter was cloned into the plasmids pSEVA237M (containing Green Fluorescent Protein, msfGFP) and pSEVA237R (containing mCherry), obtained from pSEVA collection (Durante-Rodríguez et al. 2014). The promoter was inserted by whole-plasmid PCR (primers listed in Table S1), and the PCR products were re-circularised using a KLD kit (NEB, M0554S). The resulting plasmids were transformed by electroporation into *C. testosteroni* RW31 and *P. putida* JM37, respectively, obtaining the strains *C. testosteroni* RW31-GFP and *P. putida* JM37-Cherry. Prior transformation, electrocompetent *C. testosteroni* RW31 and *P. putida* JM37 cells were prepared. Briefly, bacteria were precultured in LB O/N and inoculated at 0.1 OD_600_ in 50 mL of fresh LB. When the OD_600_ reached 0.6 - 0.8, cells were pelleted and washed twice with chilled water with 10 % of glycerol. Finally, cells were resuspended in 1 mL of water with 10 % glycerol and 100 µL aliquots were stored at - 80 ºC.

Electroporation was performed in a MicroPulser electroporator (BioRad) using 2.5 V, 200 Ω and 25 mF. Kanamycin was employed for colony selection (50 µg/mL for *P. putida* and 150 µg/mL for *C. testosteroni*).

### 2.2 Culture of Y. lipolytica HC and PET-depolymerases production

*Y. lipolytica* HC was precultured in YPD broth adjusted to pH 7 before inoculation (0.1 OD_600_) in 2 L Erlenmeyer flasks with 400 mL of Yeast Nitrogen Base minimal medium (YNB) without amino acids and nitrogen source (Difco, ref. 233520). The medium was supplemented with NH_4_Cl 0.25 g/L and glycerol 25 g/L, which corresponds to a carbon/nitrogen (C/N) ratio of ∼175. Cultures were incubated at 30 ºC and 200 rpm, collecting samples at 0, 6, 24, 48 and 72 h to monitor growth (OD_600_) and esterase activity with a *p*NPB assay (Molpeceres-García et al. 2026). *Y. lipolytica* DGA was cultured and analyzed under identical conditions as a negative control.

After 72 h, the cultures were collected and centrifuged to separate the pellet from the supernatant. The pellet was freeze-dried and stored for subsequent lipid extraction and analysis. The supernatants were stored at 4 ºC for further processing.

### 2.3 Lipid quantification and fatty acid methyl esters (FAMEs) profiling

The accumulation of lipids in *Y. lipolytica* HC was verified through staining with BODIPY (Sigma-Aldrich ref. 790389). This dye was added to culture samples at 2.5 µg/mL, observing the preparations in a DM4 B microscope (Leica) equipped with a *p*E-300 Lite LED illuminator (CoolLED) and a DFC345 FX camera (Leica).

Total lipid content was quantified by gravimetry. For lipid extraction, 200 mg of freeze-dried biomass were homogenized in 2.5 mL of 2 M HCl and heated for 1 h at 80 ºC. After cooling, lipids were extracted by adding 2 mL of methanol and then 1 mL of chloroform. Samples were vortexed for 2 min and then 1.8 mL of Milli-Q water was added. Following centrifugation, the organic phase was recovered and mixed with 2 mL of a 10 % methanol solution in chloroform. The samples were centrifuged again, recovering the organic phase and repeating the extraction process. Subsequently, chloroform was evaporated at 75 ºC in a pre-weighed tube and the lipids were measured.

To assess the lipid profile, the whole biomass was subjected to *in situ* transesterification. Around 25 mg of freeze-dried biomass were weighed, then 0.75 mg of heneicosylic acid (C21:0) was added as internal standard. Samples were then treated with 2 mL of magic methanol (methanol:hydrochloric acid:chloroform, 10:1:1), incubating at 90 ºC for 60 min. After cooling in ice, 1 mL of a 0.9 % NaCl solution and 2 mL of hexane were added, mixing in a vortex. Finally, samples were centrifuged and the organic phases, containing the corresponding FAMEs, were analysed by GC-MS (De La Torre et al. 2025).

### 2.4 Preparation of the enzymatic cocktail produced by Y. lipolytica HC with glycerol

The culture supernatants (2 L) were concentrated 40 times using 10 kDa membranes, first by tangential flow filtration (Pellicon) and then by ultrafiltration (Amicon). The final activity of the enzymatic crude was determined through the concentration process *p*NPB assay, expressed as U_*p*NPB_ /mL.

To test the activity against amPET, a piece of GoodFellow film (ES30-FM-000145) was used as substrate. The reaction mixture, composed of 30 mL of phosphate buffer 100 mM pH 8, the enzymatic cocktail at a final concentration of 40 U_pNPB_/mL, and 2.5 g of amPET, was incubated at 50 ºC and 250 rpm for five days. This assay was performed in duplicate. Four glass beads were added to each bottle to increase agitation. At final time, the samples were centrifuged and the supernatant was filtered through a 0.22 µm filter (Sartorius) for HPLC analysis of the amPET degradation products and to assess their bioavailability for *C. testosteroni* RW31 and *P. putida* JM37 (section 2.7).

### 2.5 Analysis and quantification of the degradation products from amPET

BHET, MHET and TPA were quantified in an Agilent 1200 series HPLC using a ZORBAX Eclipse Plus C18 column (retention times: 10.2 min for TPA, 11.4 min for MHET and 12.1 min for BHET) (Molpeceresc:García et al. 2026).

### 2.6 Assessment of bioavailability of the amPET hydrolysate

The degradation products obtained by enzymatic hydrolysis of amPET with the enzymatic cocktail produced in *Y. lipolytica* HC (section 2.4) were combined with MC minimal medium (Molpeceres-García et al. 2026), vitamins and trace elements, and 0.1 g/L NH_4_Cl as nitrogen source. This mixture was used to test whether *C. testosteroni* RW31 and *P. putida* JM37 could grow and produce PHAs using the molecules derived from amPET degradation as the only carbon sources. Both strains were precultured in LB and inoculated at 0.1 OD_600_ into this medium. The OD_600_ was measured after 24 h to quantify bacterial growth by turbidimetry. In addition, PHAs accumulation was assessed by staining with Nile Red at a final concentration of 5 μg/mL, as explained in Molpeceres-García et al. (2026). Granules were observed in the fluorescence microscopy mentioned above.

### 2.7 Monitoring the individual growth of P. putida GFP and C. testosteroni Cherry

*C. testosteroni* RW31-GFP and *P. putida* JM37-Cherry strains were grown individually and in co-culture in rotatory flasks in MC minimal medium supplemented with 0.1 g/L of NH_4_Cl as nitrogen source and 10 mM TPA (1.66 g/L) and 10 mM EG (0.62 g/L) as carbon sources, corresponding to C/N ratios of 36.7 and 9.2, respectively. Bacteria were precultured in LB and inoculated at OD_600_ 0.1. Their growth was monitored at 0, 6, 24 and 48 h by measuring green and red fluorescence in a SpectraMax iD3 plate reader (Molecular Devices). Absorption and emission wavelengths for GFP and Cherry were set at 480/520 nm and 575/615 nm, respectively. Growth was also monitored using the plate-counting method, as described in Molpeceres□García et al. (2026), and expressed as colony forming units per mL (CFU/mL). For that, diluted culture samples were added to two plates containing MC minimal medium with either 10 mM TPA or 10 mM EG as the carbon source. As *C. testosteroni* RW31 can assimilate TPA but not EG, while *P. putida* JM37 can assimilate EG but not TPA, this method allowed to distinguish both populations.

### 2.8 Analysis and quantification of PHAs

PHAs accumulation in the *C. testosteroni* RW31-GFP – *P. putida* JM37-Cherry consortium was visually confirmed after Nile Red and BODIPY staining employing Y3 (545/30 – 610/75 nm) and L5 (480/40 – 527/30 nm) filters (Leica), respectively, in the fluorescence microscope. Nile Red allows for PHAs assessment in *C. testosteroni* RW31-GFP, while BODIPY does for *P. putida* JM37-Cherry. In both cases, the dyes fluorescence mixes with the fluorescent protein signal and yellow granules are observed. Production of PHB and mcl-PHA was analysed and quantified by GC-MS after methanolysis of the samples as described in Molpeceres□García et al. (2026). Two analytic protocols were used, one specific for PHB, in which the retention time for C4 methyl ester was 2.4 min, and another for mcl-PHA, in which retention times were 7.0 min and 9.4 min for the C10 and C12 methyl esters, respectively.

## 3 Results and Discussion

### 3.1 Y. lipolytica HC uses glycerol to accumulate lipids and secretes active HiC and CALB

In this work we first assessed whether *Y. lipolytica* HC expressing HiC and CALB, derived from the obese strain DGA, could use glycerol to grow, accumulate lipids and secrete the recombinant enzymes at the same time. We used the background DGA strain as a control. While both strains exhibit similar growth pattern (Fig. 1), the extracellular esterase activity, measured by *p*NPB analysis, was negative in the DGA culture, and increased with the incubation time in the strain HC culture supernatants, reaching 5.2 U/mL in 72 h (Fig. 1). However, since both HiC and CALB catalyse the hydrolysis of *p*NPB, this test does not ensure that both enzymes are being produced. Hence, their presence was confirmed by SDS-PAGE. As observed in Fig. 1, the band corresponding to HiC (∼22 kDa) was much more intense than that of CALB (∼37 kDa). This could be due to a higher expression of the *hic* gene and/or to a better secretion of the mature protein, suggesting that the combination of the signal peptide SP3 with the *p*TEF promoter worked better for HiC than for CALB in this yeast chassis and using glycerol as substrate. Furthermore, the molecular mass of both proteins coincides with those of the commercial enzymes. The presence of potential *N*-glycosylation sites for HiC was discarded since the mass in the gel coincide with the theorical protein mass, and by using the NetNGlyc prediction tool (Gupta & Brunak, 2001). On the other hand, CALB has one *N*-glycosylation site at Asn74 (Uppenberg et al. 1994) and, considering that its theoretical mass according to its amino acid sequence is 33 kDa, the data suggest that *Y. lipolytica* HC produces a glycosylated form of the protein (∼11%) (Fig. 1).

**Figure 1.**
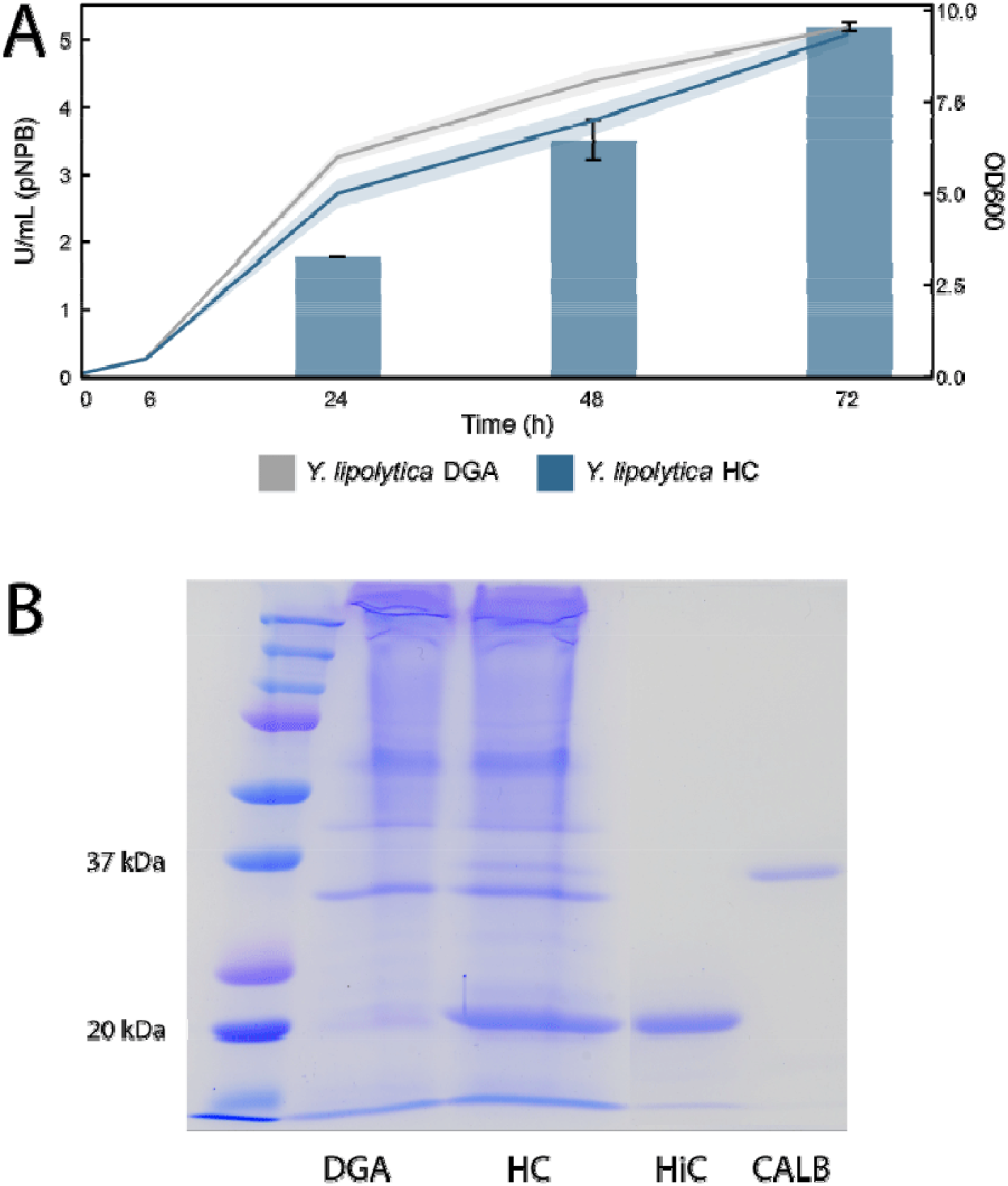
*Y. lipolytica* DGA and HC growth and production of extracellular HiC and CALB. A) Growth and esterase activity against *p*NPB. Lines represent growth (OD_600_) and bars represent extracellular esterase activity (U_*p*NPB_/mL). The strain *Y. lipolytica* DGA presented no esterase activity. Results are expressed as the mean of n = 2. Error bars and shadow areas represent standard error. B) SDS-PAGE gel showing the expression of HiC and CALB in *Y. lipolytic* HC. The first lane corresponds to molecular mass standards, and the last two lanes to commercial HiC and CALB.

According to the literature, recombinant CALB has been successfully produced in *Y. lipolytica* JMY7126, using the Lip2 signal peptide and the *p*EYK1-3AB expression system (Theron et al. 2020). Hence, different configurations could be assessed for future optimization of CALB expression in *Y. lipolytica*.

Regarding lipid production, the accumulation of intracellular TAGs was confirmed by BODIPY staining in *Y. lipolytica* HC and DGA strains (Fig. S1). Remarkably, the intracellular lipids accumulated in *Y. lipolytica* HC at 72 h constituted nearly half of the cell dry weight (CDW) in this strain (42.9 ± 1.6 %), while in DGA strains represented 39.6 ± 1.5 %. As both strains presented similar growth (Fig. 1), these results indicate that the expression of HiC and CALB no not affect lipid production. These data demonstrated the feasibility of upcycling glycerol into PET-degrading enzymes and lipids. While nitrogen limitation benefits lipid production in *Y. lipolytica* (Seip et al. 2013), it may compromise growth and PET-hydrolases production. Hence, different C/N ratios could be used to favour lipid or enzyme synthesis in *Y. lipolytica* HC, finding the most suitable trade-off for each situation.

The predominant fatty acids identified in the lipid fraction were oleic + linolenic acid (C18:1 + C18:3, 45.1 %), palmitic acid (C16:0, 25.8 %), stearic acid (C18:0, 11.9 %), linoleic acid (C18:2, 8.71 %), and palmitoleic acid (C16:1, 5.6 %), with minor amounts of others (Fig. S2). These results are similar to those reported in other studies with *Y. lipolytica* using glycerol as substrate, in which oleic acid was predominant, followed by palmitic, stearic and linoleic (Gajdoš et al., 2017; Kuttiraja et al. 2016; Papanikolaou & Aggelis, 2002). Other carbon sources and optimized fermentation systems can lead to higher accumulations with obese strains. For example, lipid production in a batch bioreactor using glucose as substrate amounted to 77 % of the CDW (Friedlander et al. 2016).

Once lipids accumulation and esterase activity were confirmed, *Y. lipolytica* HC enzymatic cocktail was tested *in vitro* against amPET employing 40 U_*p*NPB_/mL of enzyme, and yielding 7.7 ± 1.8 mM of TPA in 5 days. BHET and MHET were not detected, which is in agreement with the reported synergistic action of HiC and CALB against PET. This value is significantly higher than that previously reported by Carniel et al. (2017), who tested commercial HiC and CALB against amPET at 50 ºC and pH 8.

They obtained 2.2 mM after 14 days of reaction using a 90/10 HiC/CALB proportion and 40 mg of enzyme per g of PET, which represented a 7.7-fold increase compared to using HiC alone.

On the other hand, other studies report higher PET degradation rates by HiC using a different methodology. Kaabel et al. (2021) tested a static depolymerization in moist conditions for 7 days at 55 ºC, using 6.5 mg of commercial HiC per g of PET. Under these conditions, 20 % of a commercial semi-crystalline PET was depolymerized to TPA. This suggests that the reactions with the *Y. lipolytica* HC cocktail efficiency could be optimized by modifying the experimental settings.

### 3.2 The amPET hydrolysate can be upcycled to PHAs by a C. testosteroni RW31 – P. putida JM37 consortium

The next step consisted in evaluating whether the TPA and EG released by the enzymatic hydrolysis of amPET could be used as a metabolizable carbon source to produce PHAs in a *C. testosteroni* RW31 – *P. putida* JM37 consortium. To recover the monomers, the reaction mixture was first filtered to remove any insoluble material. Then, the culture medium was formulated by adding the components of the MC minimal medium and 0.1 g/L of NH_4_Cl as the nitrogen source. Since PHAs synthesis occurs as a response to nutritional stress, it is important to calculate the C/N ratio in the medium. However, in the co-culture each strain uses only one of the two monomers, so the C/N ratios were independently calculated for both, resulting in values of 7.1 for EG and 28.24 for TPA. *C. testosteroni* RW31 and *P. putida* JM37 were inoculated at 0.1 OD_600_ each. After 24 h, the consortium reached an OD_600_ of 0.98 ± 0.07 indicating bacterial proliferation. Additionally, the accumulation of intracellular PHAs was visualized at the microscope after staining with Nile Red (Fig. S3). These results demonstrate the feasibility of upcycling the monomers released from amPET by the enzymatic cocktail produced by *Y. lipolytica* HC.

Other strategies for PET upcycling based on two sequential steps have been documented. Welsing et al. (2025) used a recombinant strain of *Trichoderma reesei* expressing the enzyme PES-H1 to degrade amPET. After hydrolysis, a modified variant of *P. putida* upcycled the TPA and EG released into rhamnolipids, biosurfactants and biopolymers. However, the overall process presented here enables the simultaneous utilization and valorisation of two waste streams (glycerol and PET) into microbial lipids and PHAs. Also, there was no need for bacterial genetic modifications, as *C. testosteroni* RW31 and *P. putida* JM37 are natural TPA and EG assimilators and PHAs producers. While PHAs served as a rapid proof-of-concept of the biotransformation of TPA and EG into a value-added compound, both bacteria can be transformed to express heterologous metabolic pathways targeted to synthetize other value-added compounds (Molpeceres-García et al. 2026). The innovative scheme presented in this work opens the door to future consortium designs, and the distribution of biosynthetic pathways necessary to produce compounds of interest among different bacterial populations (division of labour) could increase productivity. As an example, Mehta et al. (2025) distributed the violacein biosynthesis pathway into an *E. coli* synthetic community, obtaining a 2.5-fold increased yield compared to a mono-culture expressing the whole pathway.

### 3.3 Population dynamics in the consortium C. testosteroni RW31-GFP - P. putida JM37-Cherry

Once tested the efficiency of the consortium, the bacterial strains were engineered to express fluorescent proteins that enable distinguishing each population and monitor their respective growth dynamics. *C. testosteroni* RW31was transformed with the gene encoding the GFP protein, and *P. putida* JM37 with that of the Cherry protein, obtaining the strains *C. testosteroni* RW31-GFP and *P. putida* JM37-Cherry. In both cases, fluorescence was detected in the recombinant strains. The ability of these markers to monitor population dynamics was evaluated by comparing the results with those obtained using a classical plate-counting method. In this case, two different agar plates with MC minimal medium, with either TPA or EG as carbon source, were used to differentiate each bacterium, expressing growth in CFU/mL. Both methods showed an imbalanced growth within the consortium, with predominance of *C. testosteroni* RW31-GFP over *P. putida* JM37-Cherry (Fig. 2). This behaviour was observed in a previous *C. testosteroni* RW31 – *P. putida* JM37 consortia (Molpeceresc:García et al. 2026), and could be attributed to the difficulty in coupling the growth of both bacteria given the difference in concentration of EG (10 mM, 0.62 g/L) and TPA (10 mM, 1.66 g/L) in the culture medium. In this case, we employed these equimolar concentrations because PET hydrolysis yields TPA and EG in an equimolar proportion. Moreover, we observed that *P. putida* JM37 presents a longer lag phase than *C. testosteroni* RW31 when growing in this culture medium, which could contribute to a faster growth of *Comamonas* (Molpeceres-García et al., 2026).

**Figure 2.**
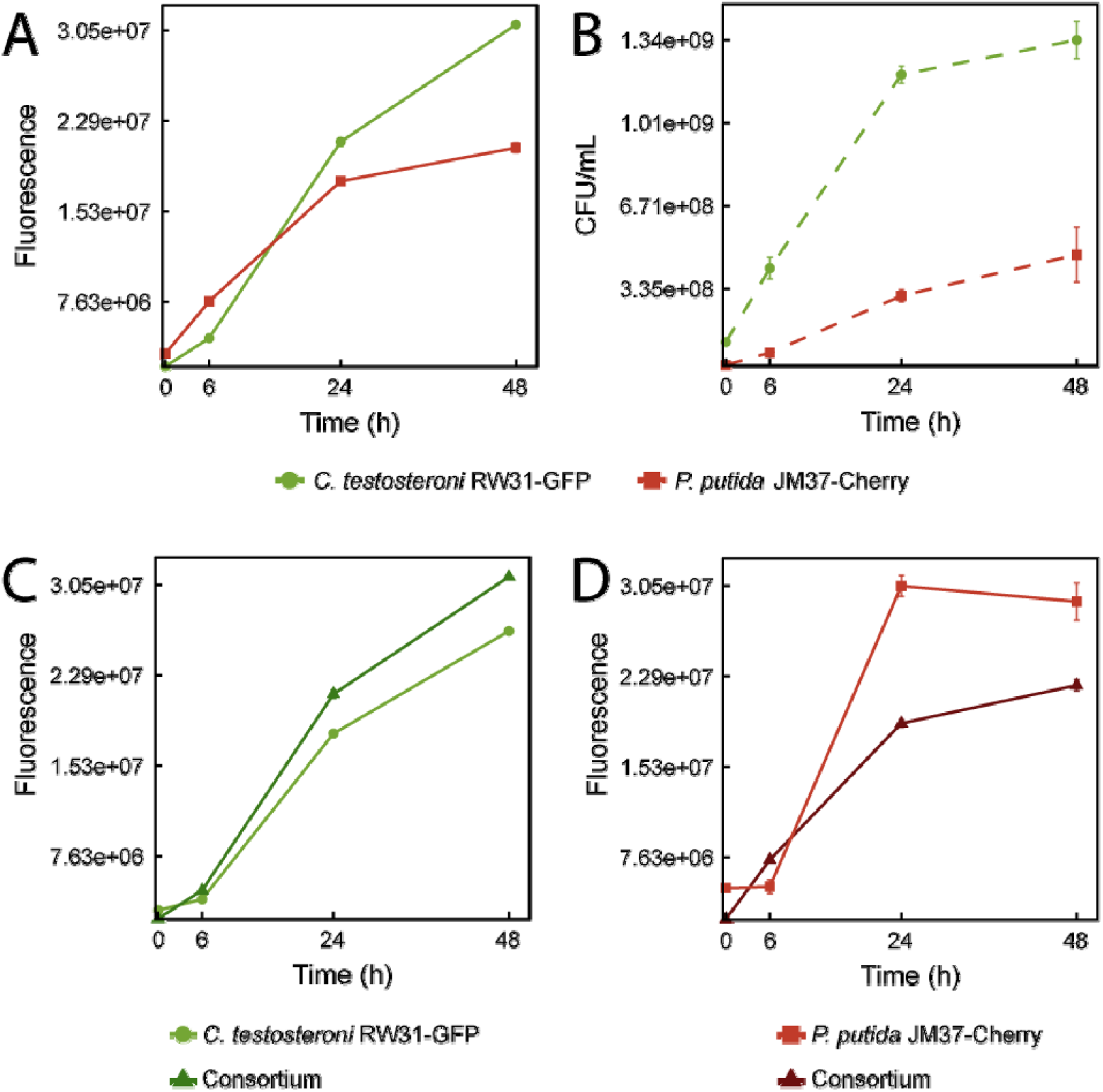
Comparation between the fluorescence and the plate-counting method for assessing the population dynamics of *C. testosteroni* RW31-GFP and *P. putida* JM37-Cherry. A) Green and red fluorescence in the consortium. B) CFU/mL of each strain in the consortium. C and D) Green and red fluorescence of monocultures of *C. testosteroni* RW31-GFP and *P. putida* JM37-Cherry compared with those detected in the coculture. Results are expressed as the mean of n = 3. Error bars represent standard error.

The metabolic pathways for degradation of each carbon source can also account for these differneces, since TPA degradation yields oxaloacetate and pyruvate, while two EG are needed for obtaining one pyruvate. The growth of *P. putida* JM37 could be enhanced adding a co-substrate, such as glucose, which *C. testosteroni* RW31 cannot assimilate (Molpeceres□García et al. 2025). In the context of PET upcycling, polycotton clothes, which are formed by PET and cotton fibres (Qiu et al. 2025), could be a good substrate, since the monomers released from its degradation (TPA, EG and glucose) could be fully upcycled by this consortium.

From a methodological perspective, some discrepancies were observed when the results obtained by fluorescence and CFU/mL were compared. While in the first case the proportions of both populations remained balanced during the first 24 h, the differences at this time were much larger according to the plate-counting method (Fig. 2). This could be explained by different yield in the expression and biosynthesis of the fluorescent proteins, even though both were cloned on plasmids with the same architecture. In the consortium, the CFU/mL of *P. putida* JM37-Cherry population was 3.9-fold lower than that of *C. testosteroni* RW31-GFP (Fig. 2). However, the fluorescent signals of both strains were similar, suggesting that *mCherry* was expressed more efficiently than *gfp*. Hence, the fluorescent intensity is not directly comparable with the population size. When the consortium is compared with the mono-cultures growing in the same coditions, the intensity of the GFP fluorescence is higher in the consortium than in the *Comamonas* single culture, while the Cherry fluorescence intensity is lower (Fig. 2) than in the *Pseudomonas* single culture. This suggests that *C. testosteroni* RW31 growth may be favoured when co-cultured with *P. putida* JM37.

Despite the differences, these methods can be complementary. While the fluorescence assay could be useful for quickly testing how different growth conditions can affect the population dynamics in a microbial consortium (Duncker et al. 2021), the plate-counting method would be more suitable for accurate growth assessment. Luu et al. (2025) reported similar issues using the Red Fluorescent Protein (RFP) as a reporter for monitoring bacterial competition in co-culture, employing two recombinant strains of *Staphylococcus aureus* and *Escherichia coli*. The authors found discrepancies in the RFP signal and the actual viable cells, as a decrease in RFP signal did not accurately correlate with a decrease in CFUs. Nonetheless, these fluorescent assays serve as semi-quantitative methods that significantly ease consortia monitoring. In this case, calculating a correction factor based on colony count results could be considered in future experiments to establish a more accurate relationship between fluorescence and growth.

### 3.4 PHAs accumulation in the consortium labelled with fluorescent proteins

Finally, the ability of the fluorescent consortium to accumulate PHAs from TPA and EG was assessed. For qualitative analysis, cells were treated with a mixed Nile Red and BODIPY staining. Nile Red allowed observing PHAs accumulation in *C. testosteroni* RW31-GFP, and BODIPY in *P. putida* JM37-Cherry. In both cases, yellow granules were observed (Fig. S4). The images suggest higher accumulation of PHB in *C. testosteroni* than of mcl-PHA in *P. putida*. Quantitative analysis of the samples after methanolysis at 24 h confirmed an accumulation of PHB of 28.4 ± 1.3 % of the cell dry weight in *Comamonas*, while in *Pseudomonas* the accumulation was not enough for quantification. However, monomers C10 and C12 from the mcl-PHA were detected in a proportion 4.7:1.

The proportion of PHB obtained agrees with that reported for a consortium formed by *C. testosteroni* RW31 and *P. putida* JM37 strains previously engineered in our group to secrete a MHETase and a PETase, respectively. Using BHET as the carbon sources, this consortium yielded 29.5% ± 0.9 % PHB/CDW, with the presence of mcl-PHA with C6, C8 and C12:1 monomers in addition to C10 and C12 (Molpeceres□García et al. 2026). Despite the only carbon source available for *P. putida* JM37 was EG in both cases, the C/N ratios were different: 9.2 in this work and 13.75 in the previous. This difference may affect mcl-PHA accumulation and monomer composition in *P. putida* (Prieto et al. 2016). However, in both studies the ratio C/N for TPA was 36.7, a value considerably higher that favour PHB accumulation by *C. testosteroni* RW31 in the consortium. Further optimization of the culture medium and the use of adequate co-substrates, such as the mentioned glucose derived from polycotton, could result in an increased population of *P. putida* JM37 and induce the accumulation mcl-PHA.

In this sense, the fine tuning of the PHB/PHAs proportion will be interesting to formulate new materials. Polymers formed by blended PHB and a low proportion of mcl-PHA have been described to improve the properties of PHB in terms of decreased crystallinity and melting temperature, increased flexibility, and resistance to elongation break (Lukasiewicz et al. 2018; Panaitescu et al. 2023; Tappel et al. 2014). Employing glycerol as carbon source, Ashby et al. (2005) co-cultured two strains of *Pseudomonas oleovorans* and *Pseudomonas corrugate* to produce PHB and mcl-PHA, reporting that modulation of the culture conditions resulted in changes of the proportion PHB/mcl-PHA that range from 34:66 to 96:4. Further studies investigating the effect of different *Comamonas-Pseudomonas* ratios on the production of PHB/mcl-PHA blends would of interest.

## 4 Conclusions

Biotechnology offers innovative solutions for the environmentally friendly valorisation of PET and other residues such as glycerol within the circular economy framework. In this study, we present a complete biological workflow for PET and glycerol upcycling. Using *Y. lipolytica* HC we proved that glycerol can be converted into a PET-active enzymatic cocktail, capable of depolymerising amorphous PET, and lipids, accumulating high intracellular proportions (up to 42.9 % of the CDW). A consortium comprising *C. testosteroni* RW31, a natural TPA assimilator, and *P. putida* JM37, a natural EG assimilator, further enables the upcycling of PET-degradation monomers into mixed PHAs, demonstrating a complete biological upcycling of PET. The recombinant strains *C. testosteroni* RW31-GFP and *P. putida* JM37-Cherry provide a straightforward method to monitor consortium dynamics, facilitating the evaluation of different configurations of the consortium. Overall, this work constitutes a proof of concept and a starting point for future strategies aimed at the biological upcycling of PET.

## Acknowledgements

This work was supported by the Spanish project MOLA (PID2024-162673NB-I00 MICIU/AEI/10.13039/501100011033 and European Union). The authors acknowledge the support toward the publication fee by the CSIC Open Access Publication Support Initiative through its Unit of Information Resources for Research (URICI).

