## Supplementary Material for "Integrated valorization of glycerol and PET into lipids and PHAs using an engineered *Yarrowia lipolytica* strain and a *Pseudomonas-Comamonas* consortium"

**Table S1.** Primers used for the cloning of a Ptrc promoter into the plasmids pSEVA237M and pSEVA237R.

| Primer ID | Sequence |
| --- | --- |
| Ptrc-Fw | TTGACAATTAATCATCCGGCTCGTATAATGGATCCTCTAGAGTCGACCTG |
| Ptrc-Rv | GAGCTCGAATTCGCGCG |

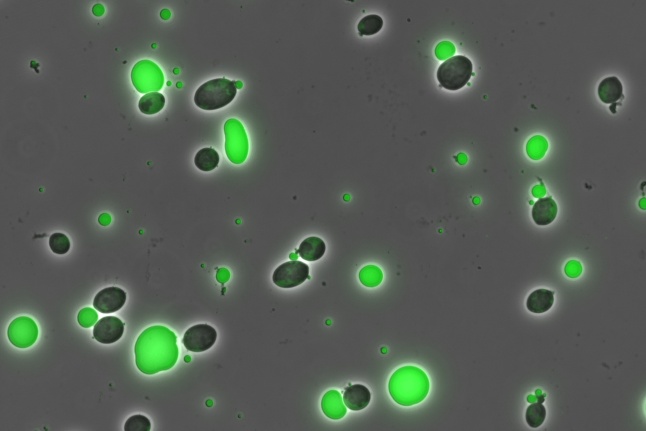

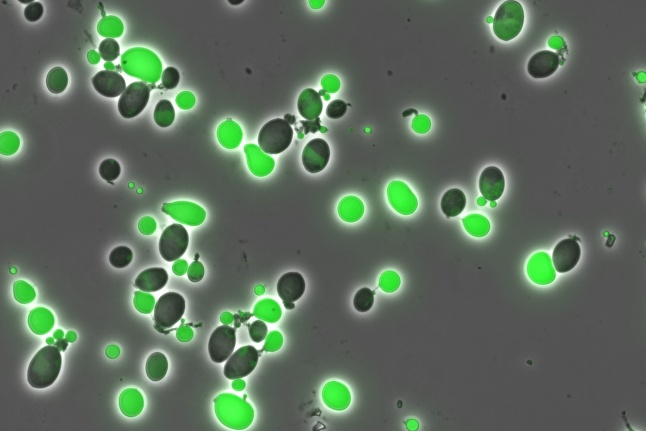

**Figure S1.** *Y. lipolytica* DGA (left) and HC (right) after 72 h growing on glycerol as the sole carbon source (C/N ratio = 175). BODIPY staining allows for the visualization of lipids accumulation.

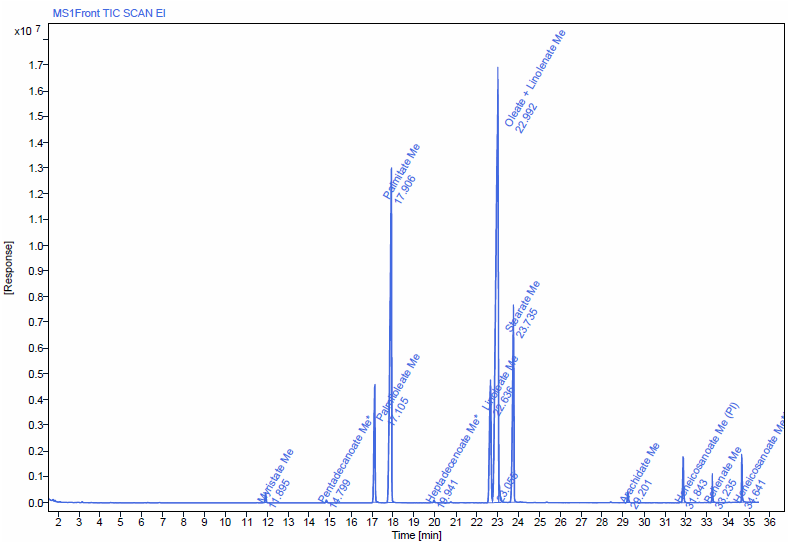

**Figure S2.** FAMES profile of *Y. lipolytica* HC growing on YNB minimal medium with glycerol as the carbon source.

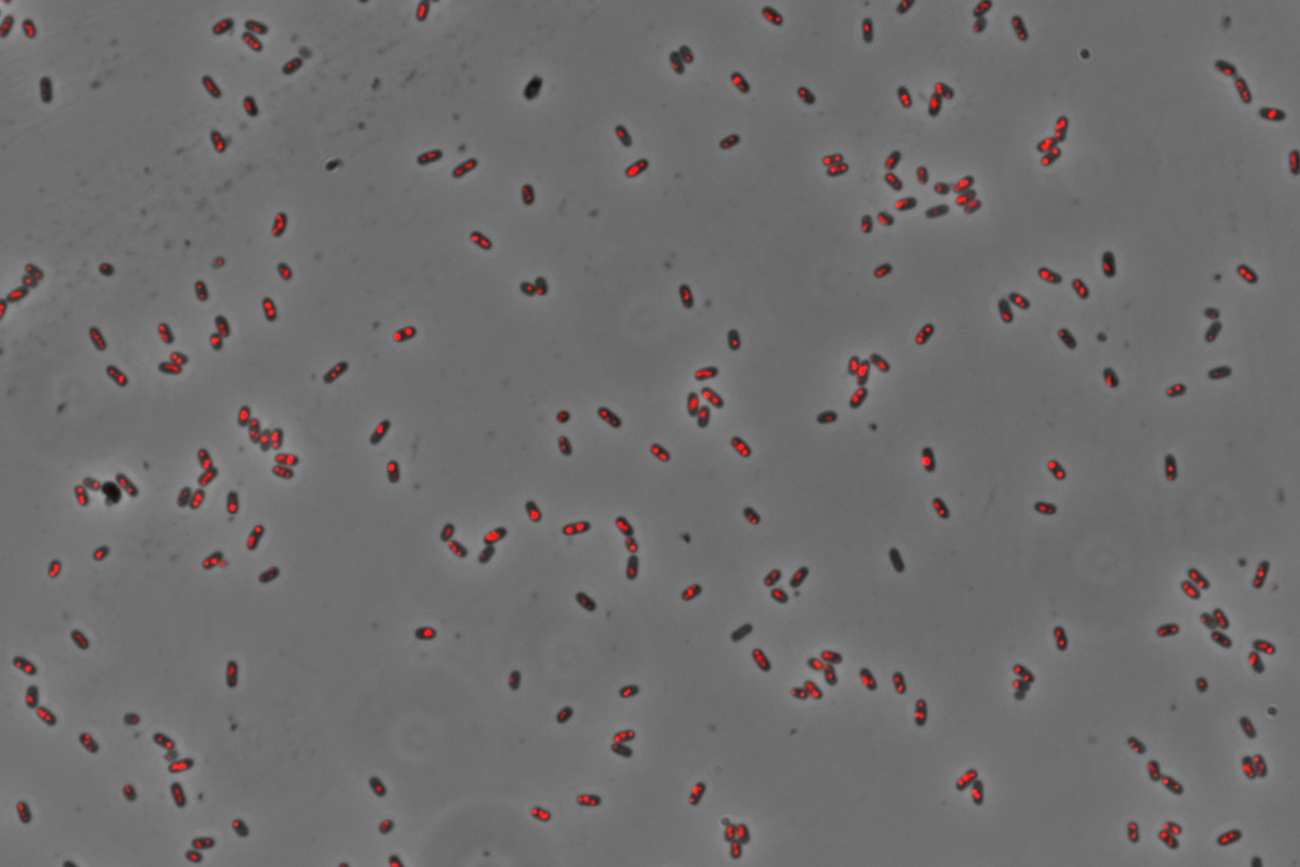

**Figure S3.** Co-culture of *C. testosteroni* RW31 - *P. putida* JM37 in MC minimal medium containing, as the carbon source, the hydrolysate resulting from enzymatic treatment of amPET with the *Y. lipolytica* HC cocktail. Nile Red staining allows visualization of PHB and mcl-PHA accumulation.

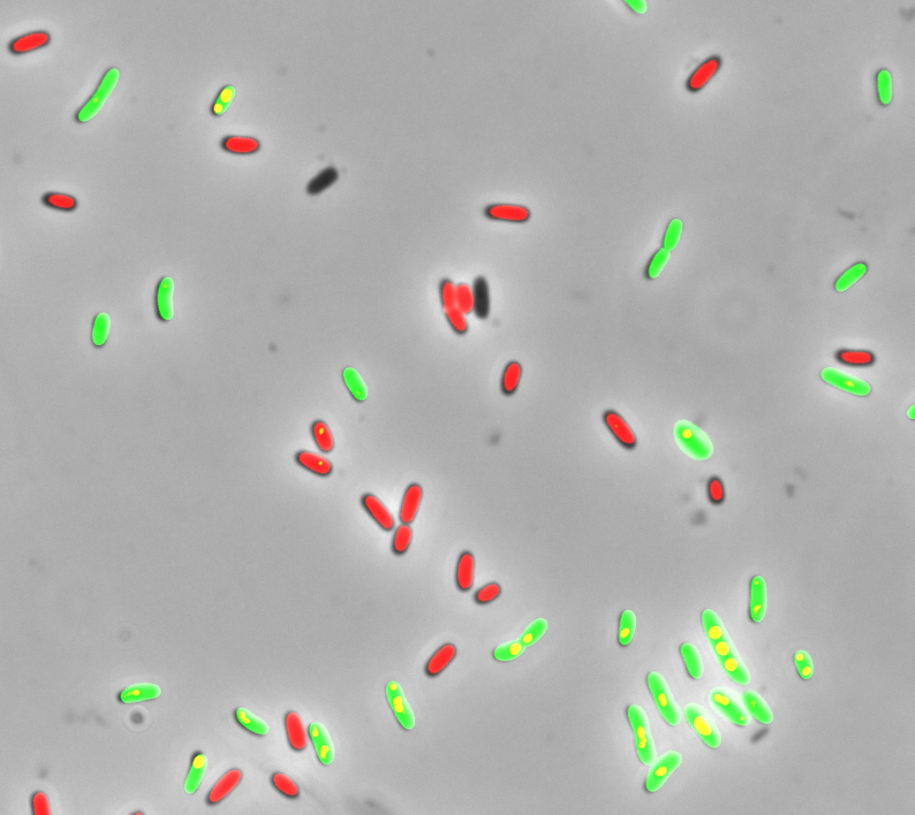

**Figure S4.** Fluorescently labelled consortium formed by *C. testosteroni* RW31-GFP and *P. putida* JM37-Cherry, stained with Nile Red and BODIPY. The mixed fluorescences and staining allows the identification of the PHB and mcl-PHA granules, as the mixed green-red signal results in yellow.
